# A Correspondence Between Normalization Strategies in Artificial and Biological Neural Networks

**DOI:** 10.1101/2020.07.17.197640

**Authors:** Yang Shen, Julia Wang, Saket Navlakha

## Abstract

A fundamental challenge at the interface of machine learning and neuroscience is to uncover computational principles that are shared between artificial and biological neural networks. In deep learning, normalization methods, such as batch normalization, weight normalization, and their many variants, help to stabilize hidden unit activity and accelerate network training, and these methods have been called one of the most important recent innovations for optimizing deep networks. In the brain, homeostatic plasticity represents a set of mechanisms that also stabilize and normalize network activity to lie within certain ranges, and these mechanisms are critical for maintaining normal brain function. In this survey, we discuss parallels between artificial and biological normalization methods at four spatial scales: normalization of a single neuron’s activity, normalization of synaptic weights of a neuron, normalization of a layer of neurons, and normalization of a network of neurons. We argue that both types of methods are functionally equivalent — i.e., they both push activation patterns of hidden units towards a homeostatic state, where all neurons are equally used — and that such representations can increase coding capacity, discrimination, and regularization. As a proof of concept, we develop a neural normalization algorithm, inspired by a phenomena called *synaptic scaling*, and show that this algorithm performs competitively against existing normalization methods on several datasets. Overall, we hope this connection will inspire machine learners in three ways: to uncover new normalization algorithms based on established neurobiological principles; to help quantify the trade-offs of different homeostatic plasticity mechanisms used in the brain; and to offer insights about how stability may not hinder, but may actually promote, plasticity.

## 1. Introduction

**S**ince the dawn of machine learning, normalization methods have been used to pre-process input data to lie on a common scale. For example, min-max normalization, unit vector normalization, z-scoring, or the like, are all well known to improve model fitting, especially when different input features have different ranges (e.g., age vs. salary). In deep learning, normalizing the input layer has also proved beneficial; for example, “whitening” input features so that they are decorrelated and have zero mean and unit variance leads to faster training and convergence [1], [2]. More recently, normalization has been extended to hidden layers of deep networks, whose activity can be viewed as inputs to a subsequent layer. This type of normalization modifies the activity of hidden units to lie within a certain range or to have a certain distribution, independent of input statistics or network parameters [3]. While the theoretical basis for why these methods improve performance has been subject to much debate — e.g., reducing co-variate shift [3], smoothening the objective landscape [4], de-coupling the length and direction of weight vectors [5], acting as a regularizer [6], [7], [8] — normalization is now a standard component of state-of-the-art architectures and has been called one of the most important recent innovations for optimizing deep networks [5].

In the brain, normalization has long been regarded as a canonical computation [9], [10] and occurs in many sensory areas, including in the auditory cortex to varying sound intensities [11]; in the olfactory system to varying odor concentrations [12]; and in the retina to varying levels of illumination and contrast [13], [14], [15]. Normalization is believed to help generate intensity-invariant representations for input stimuli, which improve discrimination and decoding that occurs downstream [9].

But beyond the sensory (input) level, there is an additional type of normalization found ubiquitously in the brain, which goes by the name of *homeostatic plasticity* [16]. Homeostasis refers to the general ability of a system to recover to some set point after being changed or perturbed [17]. A canonical example is a thermostat used to maintain an average temperature in a house. In the brain, the set point can take on different forms at different spatial scales, such as a target firing rate for an individual neuron, or a distribution of firing rates over a population of neurons. This set point is typically approached over a relatively long period of time (hours to days). The changes or perturbations occur due to other plasticity mechanisms, such as LTP or LTD, that modify synaptic weights and firing rates at much faster time scales (seconds to minutes). Thus, the challenge of homeostasis is to ensure that set points are maintained on average without “erasing” the effects of learning. This gives rise to a basic stability versus plasticity dilemma. Disruption of homeostasis mechanisms has been implicated in numerous neurological disorders [18], [19], [20], [21], [22], [23], indicating their importance for normal brain function.

In this perspective, we highlight parallels between normalization algorithms used in deep learning and homeostatic plasticity mechanisms in the brain. Identifying these parallels can serve two purposes. First, machine learners have extensive experience analyzing normalization methods and have developed a sense of how they work, why they work, and when using certain methods may be preferred over others. This experience can translate to quantitative insights about outstanding challenges in neuroscience, including the stability versus plasticity trade-off, the roles of different homeostasis mechanisms used across space and time, and whether there are parameters critical for maintaining homeostatic function that have been missed experimentally. Second, there are many normalization techniques used in the brain that have not, to our knowledge, been deeply explored in machine learning. This represents an opportunity for neuroscientists to propose new normalization algorithms from observed phenomena or established principles [24] or to provide new perspectives on why existing normalization schemes used in deep networks work so well in practice.

## 2. Normalization methods across four spatial scales

We begin by describing artificial and neural normalization strategies that occur across four spatial scales (Figure 1, Table 1): normalization of a single neuron’s activity via intrinsic neural properties; normalization of synaptic weights of a neuron; normalization of a layer of neurons; and normalization of an entire network of neurons.

**Fig. 1:**
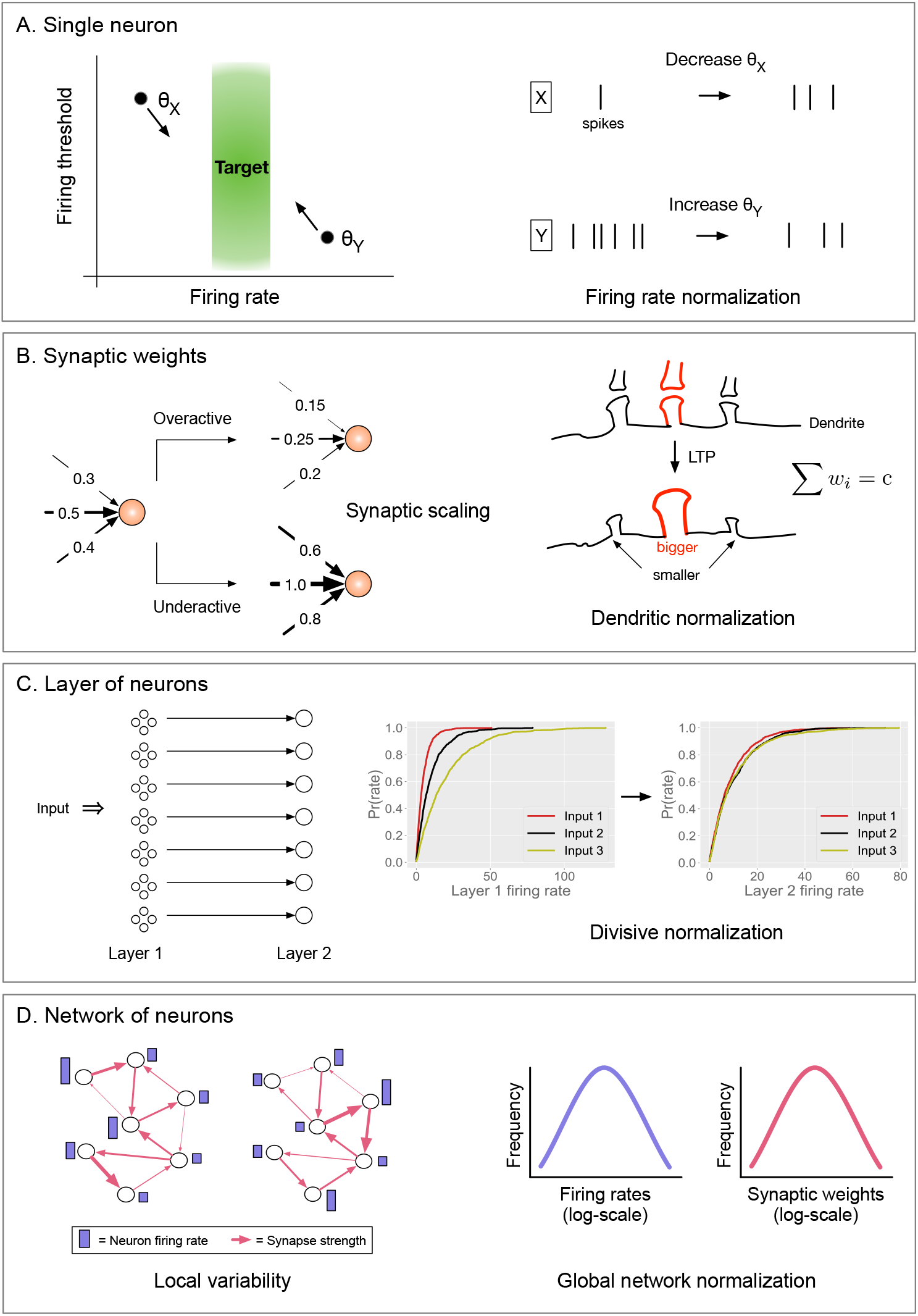
Neural homeostatic plasticity mechanisms across four spatial scales. A) Normalization of a single neuron’s activity. Left: Neuron *X* has a relatively low firing rate and a high firing threshold, *θ*_*X*_, and vice-versa for neuron *Y*. Right: Both neurons can be brought closer to their target firing rate by decreasing *θ*_*X*_ and increasing *θ*_*Y*_. B) Normalization of synaptic weights. Left (synaptic scaling): If a neuron is firing above its target rate, its synapses are multiplicatively decreased, and vice-versa if the neuron is firing below its target rate. Right (dendritic normalization): If a synapse size increases due to strong LTP, its neighboring synapses decrease their size. C) Normalization of a layer of neurons. Left: Two layers of neurons with feed-forward connections, and other feed-back inhibitory connections (not shown). Right: The cumulative distribution of firing rates for neurons in the first layer is exponential with a different mean for different inputs. The activity of neurons in the second layer are normalized such that the means of the three exponentials are approximately the same. D) Left: Example of a neural circuit with the same units and connections, but different activity levels for neurons (purple bars) and different weights (pink arrow thickness) under two different conditions. Right: Despite local variability, the global distributions of firing rates and synaptic weights for the network remains stable (log-normally distributed) under both conditions.

**TABLE 1:**
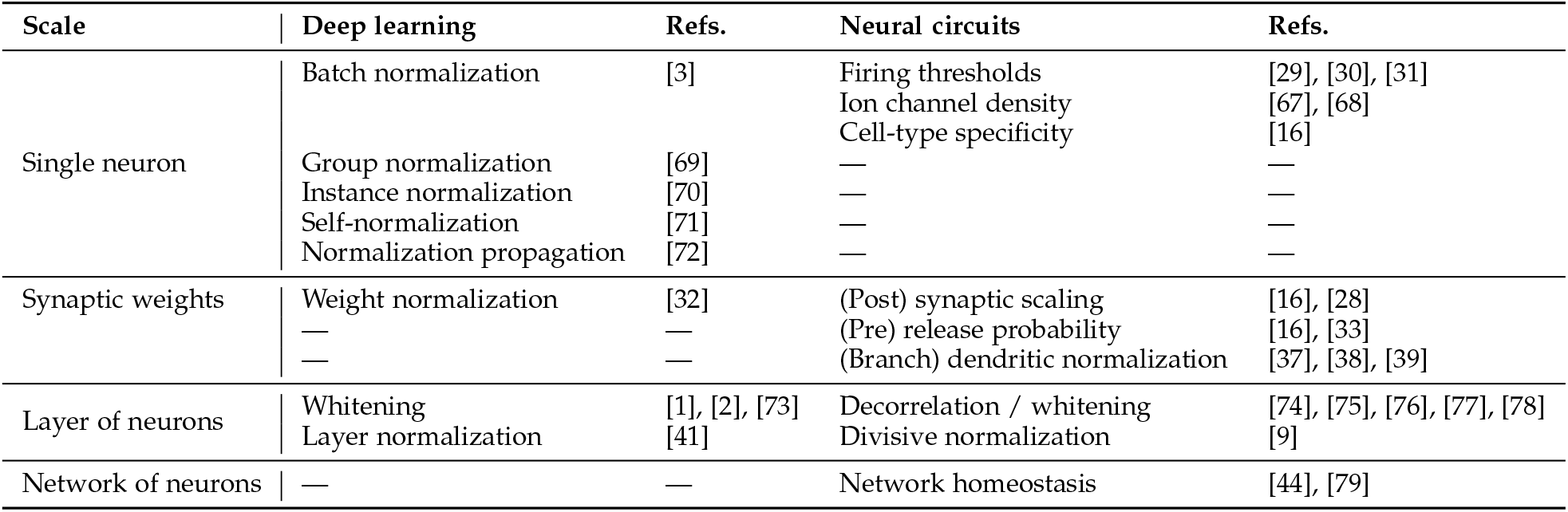
Correspondences between normalization mechanisms in artificial and biological neural networks across four spatial scales. See also reviews: [34], [64], [65], [66].

### 2.1 Normalization of a single neuron’s activity

Here, we focus on normalization methods that directly modify the activity level of a neuron via intrinsic mechanisms.

In deep learning, the current most popular form of single neuron normalization is called *batch normalization* [3]. It has long been known that z-scoring the input layer — i.e., shifting and scaling the inputs to have zero mean and unit variance — speeds up network training [1]. Batch normalizaton essentially applies this idea to each hidden layer by ensuring that, for every batch of training examples, the activation of a hidden unit over the batch has zero mean and unit variance.

Mathematically, let {*z*_1_, *z*_2_, …, *z_B_*} be the activations of hidden unit *z* for each of the *i* = 1 : *B* inputs in a training batch. Let *μ_B_* and 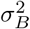 be the mean and variance of all *z_i_*’s, respectively. Then, the batch-normalized activation of *z* for the *i*^th^ input is:

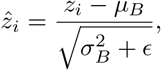

where *ϵ* is a small constant.

In practice, the effect of this simple transformation is profound: it leads to significantly faster convergence (larger learning rates) and improved stability (less sensitivity to parameter initialization and learning rate) [3], [4], [25], [26]. Numerous extensions of this method have since been proposed with various tweaks and perks on a similar underlying idea (Table 1).

In the brain, normalizing the activity of a neuron has long been appreciated as an important stabilizing mechanism [27]. For example, if neuron *u* drives neuron *v* to fire, the synapse between them may get strengthened by Hebbian plasticity. Then, the next time *u* fires, it is even more likely that *v* fires, and this positive feedback loop can lead to excessive activity. Similarly, if the synapse undergoes depression, then it is less likely for *v* to fire in the future, and this negative feedback can lead to insufficient activity. The job of homeostasis is to prevent neurons from being both over-utilized (hyperactive) and under-utilized (hypoactive) [28].

Modifying a neuron’s excitability — e.g., its firing threshold or bias — represents one intrinsic neural mechanism used to achieve homeostasis [29], [30], [31]. The idea is simple (Figure 1A); each neuron has an approximate target firing rate at which it prefers to fire. A neuron with sustained activity above its target will increase its firing threshold, such that it becomes harder to fire, and likewise, a neuron with depressed activity below its target will decrease its firing threshold, thus becoming more sensitive to future inputs. The net effect of these modifications is that the neuron hovers around its target firing rate, on average over time. Several parameters are involved in this process, such as the rate at which thresholds are adjusted, which affects how quickly homeostasis is approached, and the value of the target itself, which may be cell-type specific. Other intrinsic mechanisms, such as modifying ion channel density, can also be used to intrinsically regulate firing rates (Figure 1).

Both of these methods are unsupervised; they adjust the activity of a neuron to lie within a preferred, narrow range with respect to recently observed data.

### 2.2 Normalization of synaptic weights

Here, we focus on normalization methods that indirectly modify the activity of a neuron by changing its weights.

In deep learning, one popular way to normalize the inputs to a unit (post-synaptically) is called *weight normalization* [32]. The main idea is to re-parameterize the conventional weight vector **w** of a unit into two components:

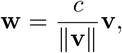

where *c* is a scalar and **v** is a parameter vector, both of which are learned. This transformation fixes the length (Euclidean norm) of the weight vector, such that ||**w**|| = *c*, for any **v**. Backpropagation is then applied to *c* and **v**, instead of to **w**. Thus, the length of the weight vector (*c*) is de-coupled from the direction of the weight vector (**v**/||**v**||). Such “length-direction” decoupling leads to faster learning and exponential convergence in some cases [5].

In the brain, the most well-studied type of weight normalization is called *synaptic scaling* [28] (Figure 1B, Left). If a neuron is on average firing above its target firing rate, then all of its incoming excitatory synapses are downscaled (i.e., multiplied by some factor, 0 < *α* < 1), to reduce its future activity. Similarly, if a neuron is firing far below its target, then all its excitatory synapses are upscaled (*α* > 1); in other words, prolonged inactivity leads to an *increase* in synaptic size [33]. These rules may seem counter-intuitive, but remember that these changes are happening over longer time scales than the changes caused by standard plasticity mechanisms. Indeed, it is hypothesized that one way to resolve the plasticity versus stability dilemma is to temporally segregate Hebbian and homeostatic plasticity so that they do not interfere [34]. This could be done, for example, by activating synaptic scaling during sleep [35], [36].

Interestingly, synapse sizes are scaled on a per neuron basis using a multiplicative update rule (Figure 1B, Left). For example, if a neuron has four incoming synapses with weights 1.0, 0.8, 0.6, and 0.2, and if the neuron is firing above its target rate, then the new weights would be downscaled to 0.5, 0.4, 0.3, and 0.1, assuming a multiplicative factor of *α* = 1/2. Critically, multiplicative updates ensure that the relative strengths of synapses are preserved, which is believed to help maintain synapse-specificity of the neuron’s response caused by learning. The value of the multiplicative factor need not be constant, and could depend, for example, on how far away the neuron is from reaching its target rate. Thus, synaptic scaling keeps the firing rate of a neuron within a range while preserving the relative strength between synapses.

Another form of weight normalization in the brain is called *dendritic normalization* [37], [38], [39], which occurs locally on individual branches of a neuron’s dendritic arbor (Figure 1B, Right). The idea is that if one synapse gets strengthened, then its neighboring synapses on the arbor compensate by weakening. This process is homeostatic because the total strength of all synapses along a local part of the arbor remains approximately constant. This process could be mediated by a shared resource, for example, a fixed number of post-synaptic neurotransmitter receptors available amongst neighboring synapses [40]. Computationally, this process creates sharper boundaries between spatially adjacent synapses receiving similar inputs, which could enhance discrimination and contrast.

### 2.3 Normalization of a layer of neurons

Here, we focus on normalization schemes that modify the activity of an entire layer of neurons, as opposed to just a single neuron’s activity.

In deep learning, *layer normalization* [41] was recently proposed to overcome several drawbacks of batch normalization. In batch normalization, the mean and variance statistics of each neuron’s activity is computed across a batch of training examples, and then each neuron is normalized with respect to its own statistics over the batch. In layer normalization, the mean and variance is instead computed over an entire layer of neurons for each training example, and then each neuron in the layer is normalized by the same mean and variance. Thus, layer normalization can be used online (i.e., batch size of one), which makes it more amenable to training recurrent neural networks [41].

In the brain, layer-wise normalization has most prominently been observed in sensory systems (Figure 1C, Left). For example, in the fruit fly olfactory system, the first layer of (receptor) neurons encode odors via a combinatorial code, in which, for any individual odor, most neurons respond at a low rate, and very few neurons respond at a high rate [42]. Specifically, the distribution of firing rates over all receptor neurons is exponential with a mean that depends on the concentration of the odor (higher concentration → higher mean). In the second layer of the circuit, projection neurons receive odor excitation from receptor neurons, as well as inhibition from lateral inhibitory neurons [12]. The result is that the concentration-dependence is largely removed; i.e., the distribution of firing rates for projection neurons follows an exponential distribution with approximately the same mean, for all odors and all odor concentrations [43] (Figure 1C, Right). Thus, while an individual neuron’s firing rate can change depending on the odor, the distribution of firing rates over all neuron’s remains nearly the same for any odor. This process is dubbed *divisive normalization* and is believed to help fruit flies identify odors independent of the odor’s concentration. Divisive normalization has also been studied in the visual system, for example, light adaptation in the retina, or contrast adjustment in the visual cortex [9].

Overall, layer normalization helps maintain some invariant response property of a layer of neurons by dividing the responses of individual neurons by a factor that relates to the summed activity of all the neurons in the layer. These normalizations can be considered “homeostatic” because they preserve, for any input, properties of a distribution of firing rates (e.g., the mean or variance). In the brain, other non-linear transformations are also used alongside these transformations, for example, to adjust saturation rates of individual neurons and to amplify signals prior to normalization [9].

### 2.4 Normalization of a network of neurons

In the brain, recent work has challenged the conventional view that homeostasis applies at the level of a single neuron or a strict layer of neurons, and have instead attributed homeostasis properties to a broader network of neurons. In one experiment, the firing rates of individual neurons in a hippocampal network were monitored for two days after applying baclofen, a chemical agent that suppresses neural activity. After two days, the distribution of firing rates over the population was compared to the distribution of firing rates for a control group of neurons that received no baclofen. Strikingly, both were approximated by the same log-normal distribution. Moreover, the firing rates of many individual neurons, in both conditions, significantly changed from day 0 to day 2 [44], [45] (Figure 1D), suggesting that homeostasis may not strictly apply at the level of an individual neuron but is rather maintained at the population level. Similar observations have been made in the stomatogastric ganglion of crabs and lobsters, where rhythmic bursting is robustly maintained despite many perturbations to the circuit [46]. This remains a beautiful yet mysterious property of network stability implemented by neural circuits, and the mechanisms driving this level of network regulation remain poorly understood [47].

In deep learning, we are not aware of a normalization strategy that is applied across an entire network of units, or even across a population of units beyond a single layer. Network homeostasis could in principle be an emergent property from local homeostasis rules implemented by individual units, or could be a global constraint intrinsically enforced by some unknown mechanism. Either way, we hypothesize that network homeostasis may be attractive in deep networks because it allows for more flexible local representations while still providing stability at the network level.

## 3. The benefits of homeostasis (load balancing)

In computer science, the term “load balancing” means to distribute a data processing load over a set of computing units [48]. Typically, the goal is to distribute this load evenly to maximize efficiency and reduce the amount of time that units are idle (e.g., for servers handling traffic on the Internet). For neural networks, we define load balancing based on how frequently a set of neurons are activated, and how similar their mean activation levels are, on average. Why might load balancing in neural networks be attractive computationally? Three reasons come to mind:

First, load balancing increases the coding capacity of the network; i.e., the number of unique stimuli that can be represented using a fixed number of resources (neurons). Suppose that under standard training, a certain fraction (say, 50%) of the hidden units are just not used; that is, they are never, or rarely ever, activated. This wasted capacity would reduce the number of possible patterns the network could represent and would introduce unnecessary parameters that can prolong training. Load balancing of neurons could avoid these problems by pressing more hidden units into service. In the brain, equal utilization of neurons also promotes distributed representations, in which each stimuli is represented by many neurons, and each neuron participates in the representation of many stimuli (often called a combinatorial code [42], [49]). This property is particularly attractive when such representations are formed independent of input statistics or structure.

Second, load balancing can improve fine-grained discrimination. Suppose there are two hidden units that are similarly activated for the same input stimuli (e.g., images of dogs). The training process could just choose one of them and turn off the other. But if both units are used, then the door remains open for future fine-grained discrimination; e.g., discriminating between subclasses of dogs, such as chihuahuas and labradoodles. In general, if more nodes are used to represent a stimulus, then the nodes may better preserve finer details of the pattern, which can serve later as the basis for discrimination, if necessary. Relatedly, if a neuron has a sigmoidal activation function, normalization keeps the neuron in its non-saturated regime. This is believed to help the neuron be maximally informative and discriminative [50], [51], [52], [53], [54].

Third, load balancing can serve as a regularizer, which is commonly used in deep networks to constrain the magnitude of weights or the activity levels of units. Regularizers typically improve generalization and reduce over-fitting [55], and can be specified explicitly or implicitly [56]. There are many forms of regularization used in deep learning; for example, Dropout [57], in which a random fraction of the neurons is set inactive during training; or weight regularization, in which *ℓ*_1_ or *ℓ*_2_ penalties are applied to the loss function to limit how large weight vectors become [58], [59]. Although regularization is a powerful tool to build robust models, regularization alone is not guaranteed to generate homeostatic representations.

## 4. Empirically testing the benefits of homeostasis

The empirical results in this section serve two purposes. The first is to show that two popular normalization methods generate homeostatic representations and demonstrate the three benefits of homeostasis discussed in the previous section. The second is to show that a method inspired by synaptic scaling also does the same and performs competitively against existing normalization methods. These results are not meant to represent a full-fledged comparison between normalization methods across multiple architectures, datasets, or hyper-parameter settings. Rather, these results are simply meant to demonstrate a proof-of-concept of the bi-directional perspective argued here.

### Experimental setup

For our basic architecture, we used the original LeNet5 [60] with two convolutional layers and three fully connected layers with ReLU activation functions. Like neurons, ReLU units include a firing threshold; only nodes with a value greater than a threshold can fire.

We experimented with two datasets. The first is CIFAR-10, a standard benchmark for classification tasks, which contains 60,000 color images, each of size 32 × 32, and each belonging to one of 10 classes (airplanes, cats, trucks, etc.). The second dataset is SVHN (Street House View Numbers), which contains 73,257 color images, each of size 32 × 32, and each belonging to one of 10 classes (digits from 0–9). SVHN is analogous to MNIST but is more difficult to classify because it includes house numbers in natural scene images taken from a street view.

Each normalization method is applied to every layer, except the input and output layers, with all affine parameters trainable. For each dataset, all methods used Adam optimization using PyTorch with default parameters. Additional hyper-parameters were fixed for each dataset: CIFAR-10 (batch size of 32, learning rate of 0.003, train for about 45,000 iterations), SVHN (batch size of 256, learning rate of 0.01, train for about 8,000 iterations). Batch statistics are calculated using training data during training and testing data during testing. Table 2 provides the equations for each normalization algorithm.

**TABLE 2:**
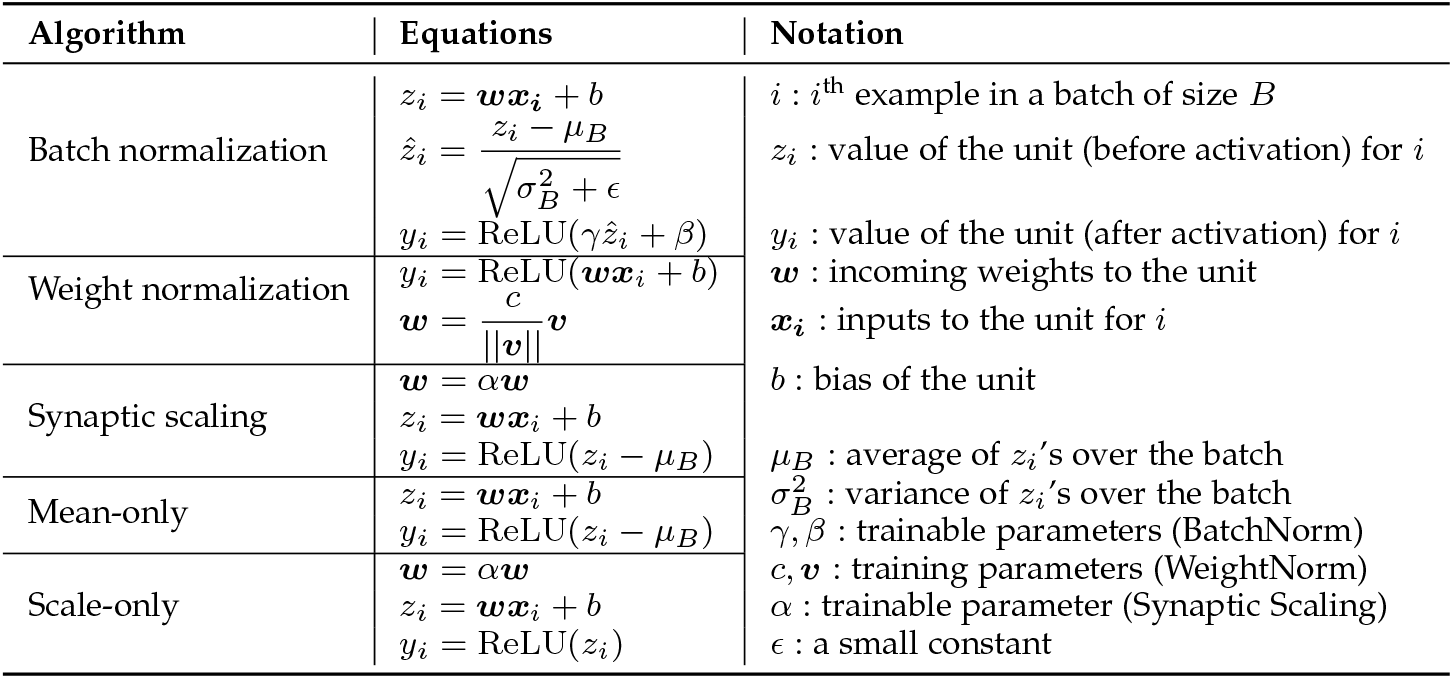
Normalization algorithms. All equations show the forward-pass update equations for a single hidden unit. For Weight normalization, backpropagation is performed on *c* and ***v***, instead of ***w***.

The three benefits of homeostasis are measured as follows: 1) Coding capacity: the information entropy of the binarized activation values over hidden units. Entropy is highest when the probability that a random unit is activated (i.e., it outputs a value > 0) for an input is 50%. 2) Discrimination: the classification accuracy on the test set. 3) Regularization: the range of the response magnitudes of hidden units; a narrower range implies better regularization.

### A synaptic-scaling-inspired normalization algorithm

Of the many normalization methods discussed above, we choose to model synaptic scaling because it is one of the most well studied and widely-observed mechanisms across brain regions and species.

We propose a simplified model of synaptic scaling that captures two keys aspects of the underlying biology: multiplicative scaling of synaptic weights, and constraining a node to be activated around a target activation probability on average. In the first step, the incoming weight vector ***w*** for a hidden unit is multiplied by a factor *α*, i.e., ***w*** = *α**w***. Each hidden unit has its own *α* value, which is made learnable during training. The *α* values are initialized to 1. In the second step, for each hidden unit, we subtract its mean activation (over a batch) from its actual activation for each input in the batch. This process ensures that each unit has a mean activation (before ReLU) of 0 and hence, a probability of activation (output value > 0) of around 50%, and thus resembles the biological observation that no neuron is over- or under- utilized. This step is also the same as mean-only batch normalization [32]. One advantage of this synaptic scaling model compared to batch normalization is that it removes the division by the variance term, which can lead to exploding gradients when the variance is close to zero.

An “ideal” model of synaptic scaling might only multiplicatively scale the weights of a hidden unit such that a given target activation probability is achieved on average. Instead, we first scale the weights by a learnable parameter (*α*), which allows the network to learn the optimal range of activation values for the unit, and we then constrain the unit to hit its target in step two. Similarly, batch normalization does not simply use z-scored activation values for each hidden unit (Table 2), but rather includes two learnable parameters (*γ, β*) per unit to shift and scale its normalized activation. In both cases, this flexibility likely increases the representation power of the network [3].

Mathematically, for each hidden unit, the forward-pass operations for synaptic scaling are:

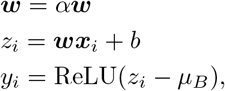

where the subscript *i* indicates the *i*^th^ example in a batch of size *B* (*i* = 1 : *B*); ***w***, ***x_i_***, *b*, *z_i_*, *y_i_* are the incoming weights to the hidden unit, the inputs for the *i*^th^ example from the previous layer, the bias of the unit, the value of the unit before activation, and the output of the unit, respectively; *μ_B_* is the average of all *z_i_*’s over a batch.

To explore how the two steps independently affect classification performance, we tested each of them without the other. We call these models “Mean-only” and “Scale-only”, respectively (Table 2).

### 4.1 Existing normalization methods generate homeostatic representations

First, we confirmed that two state-of-the-art normalization methods — batch normalization (BatchNorm) and weight normalization (WeightNorm) — improve discrimination (i.e., classification accuracy) on CIFAR-10: from 59.3 ± 1.4% for the original version of LeNet5 without normalization (Vanilla) to 63.8 ± 0.9% (WeightNorm) and 65.8 ± 0.5% (BatchNorm) (Figure 2A). Normalized networks also learned faster; they required fewer training iterations to achieve high accuracy.

**Fig. 2:**
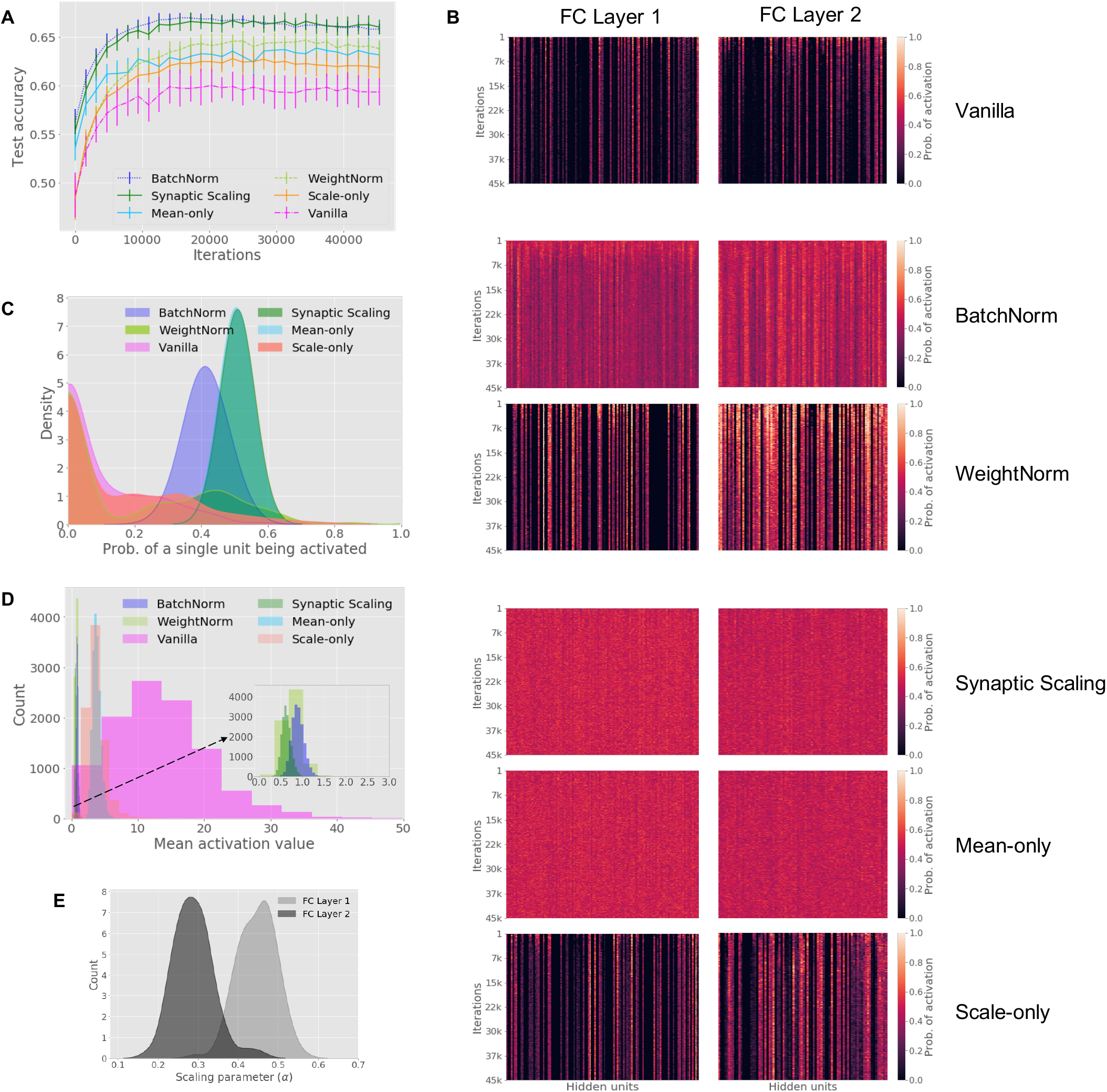
[CIFAR-10] Normalization increases performance and drives neural networks towards a “homeostatic” state. A) Test accuracy (*y*-axis) versus training iteration (*x*-axis). Error bars show standard deviation over 10 random initializations. BatchNorm and Synaptic Scaling achieve higher accuracy at the beginning and at the end of training compared to all other methods, including Vanilla. B) The probability of each hidden unit (columns) being activated over all inputs in a batch, computed on every 100^th^ training iteration (rows). Heatmaps are shown for hidden units in both fully-connected (FC) layers. C) Distribution of the probabilities that each unit in the first FC layer is activated per input. D) Histogram of the mean activation values for hidden units in the first FC layer, calculated using the test dataset. E) Distribution of the trained *α* parameters for Synaptic Scaling, for each FC layer.

Second, we show that BatchNorm and WeightNorm have higher coding capacity; i.e., units are relatively equally utilized, each with an activation probability close to 0.50. Figure 2B shows that hidden units in normalized networks had more similar activation probabilities than in Vanilla: the coefficients of variation of activation probabilities across hidden units were 0.20 (BatchNorm) and 1.38 (WeightNorm) compared to 1.65 (Vanilla). Further, individual units in networks with BatchNorm and WeightNorm had activation probabilities closer to 0.50 compared to Vanilla. For example, in the first fully-connected layer, units in BatchNorm and WeightNorm had activation probabilities of 0.41 ± 0.05 and 0.17 ± 0.23, respectively, compared to Vanilla (0.10 ± 0.15) (Figure 2C). For BatchNorm, the probability of activation forms a Gaussian-like distribution, whereas for Vanilla, the distribution forms a huge peak at 0.0 with a long right tail, indicating that many units in Vanilla never get activated, while a few units are “over used”. The distribution of WeightNorm lies between Vanilla and BatchNorm.

Third, for regularization, Figure 2D shows that when active, the values of hidden units have a narrower distribution when using normalization compared to without normalization; the coefficients of variation of activation values across hidden units were 0.16 (BatchNorm) and 0.30 (WeightNorm) compared to 0.55 (Vanilla). The average output value for hidden units was also significantly reduced in BatchNorm (0.89 ± 0.14) and WeightNorm (0.76 ± 0.23) compared to Vanilla (13.36 ± 7.36). By reducing the activation values of hidden units and confining them to a narrower range, BatchNorm and WeightNorm demonstrate regularization.

Overall, BatchNorm does a better job of pushing the network towards a homeostatic state compared to WeightNorm and that may contribute to why BatchNorm also achieves better classification accuracy than WeightNorm.

### 4.2 Synaptic scaling performs load balancing and obtains competitive performance

We next tested the Synaptic Scaling method and found that its classification accuracy (66.0 ± 0.7%) was very similar to BatchNorm (65.8 ± 0.5%) on CIFAR-10 (Figure 2A). In contrast, Mean-only and Scale-only performed worse than Synaptic Scaling, suggesting that both steps — multiplicative scaling of synapses and setting target activation probabilities — are better when combined.

Synaptic Scaling also has a coding capacity that is on par or even slightly better than BatchNorm. Figure 2B shows that each hidden unit had a very similar probability of being activated; coefficient of variation of 0.11 for Synaptic Scaling and 0.20 for BatchNorm, compared to 1.65 for Vanilla. Synaptic Scaling activated hidden units with a probability of 0.51 ± 0.02, which is slightly higher than BatchNorm (0.41 ± 0.05) and much higher than Vanilla (0.10 ± 0.15) (Figure 2C).

Finally, for regularization, Figure 2D shows that the activation values across hidden units were similar after normalization; coefficient of variation of 0.17 for Synaptic Scaling and 0.16 for BatchNorm, compared to 0.55 for Vanilla. The average output value for hidden units was also reduced in Synaptic Scaling and BatchNorm (0.63 ± 0.11 versus 0.89 ± 0.14, respectively) compared to Vanilla (13.36 ± 7.36).

Interestingly, the learned *α* parameters for Synaptic Scaling are all positive, meaning no weights flipped sign during training, and all the *α* < 1, meaning the weights are all scaled down (Figure 2E). We did not set any upper or lower bounds on *α*, and the fact that the learned values stay within [0, 1] indicates that down-scaling of weights, which in turn reduce activation values, may generally be beneficial for this classification task.

#### Validation on a second dataset

To ensure these results were not specific to one dataset, we ran all methods on a second dataset (SVHN) and found similar trends (Figure 3). To summarize, Synaptic Scaling and BatchNorm improve classification accuracy (Figure 3A), coding capacity (Figure 3B-C), and regularization (Figure 3D), compared to all other methods.

**Fig. 3:**
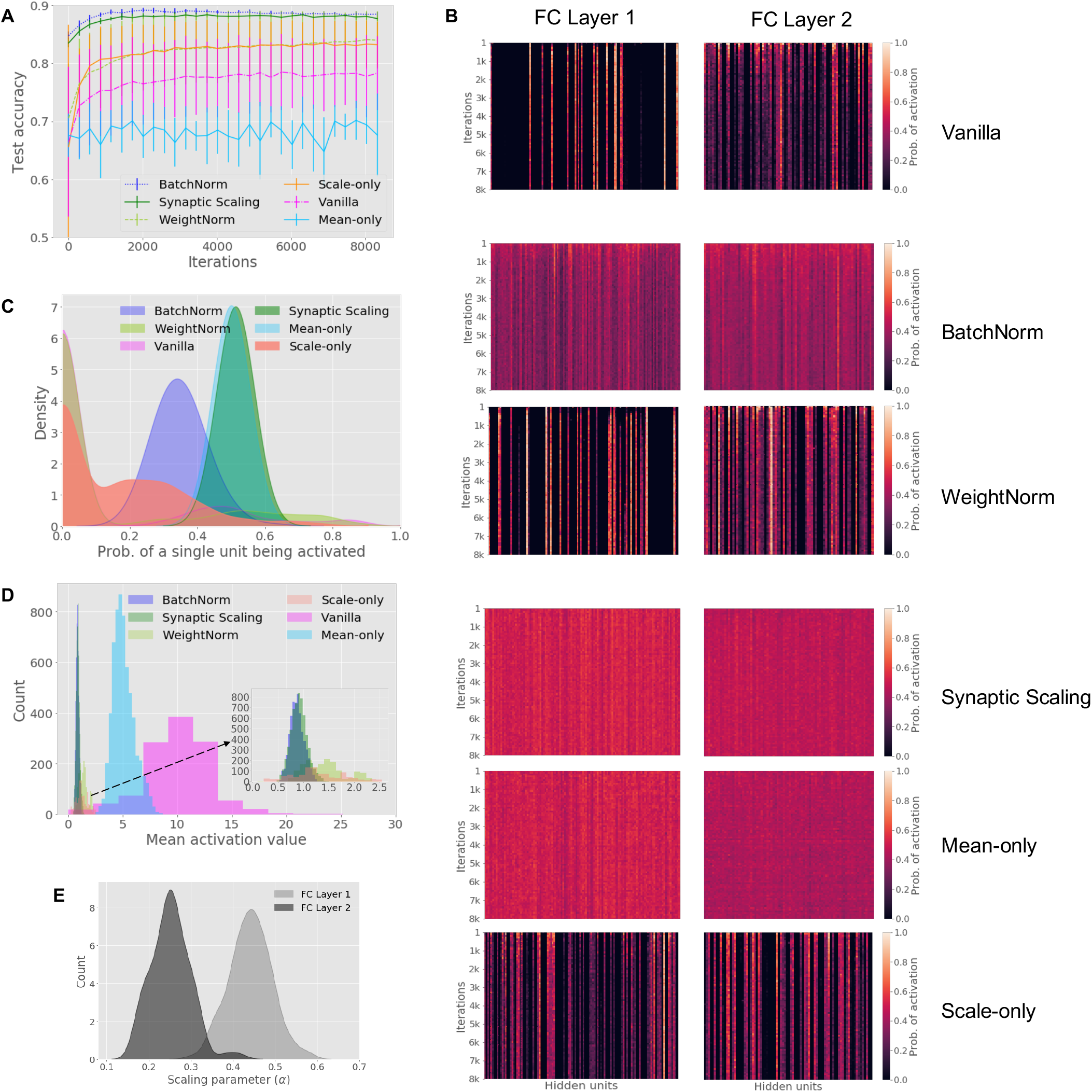
[SVHN] Similar benefits of normalization on a second dataset. Synaptic Scaling and BatchNorm have the highest classification accuracy (A), increase coding capacity (B,C: probability of each hidden unit being activated), and increase regularization (D: mean activation values for hidden units). See Figure 2 caption for detailed panel descriptions.

## 5. Discussion

We showed that widely used normalization methods in deep learning are functionally equivalent to homeostatic plasticity mechanisms in the brain. While the implementation details vary, both ensure that the activity of a neuron is centered around some fixed value or lies within some fixed distribution, and both are temporally local in the sense that changes only depend on recent behavior (recent firing rate or recent data observed). In summary, both attempt to stabilize and bound neural activity in an unsupervised manner, and both are critical for efficient learning.

We showed that two state-of-the-art normalization methods (BatchNorm and WeightNorm), as well as a new normalization algorithm inspired by synaptic scaling, demonstrate the three benefits of load balancing: compared to Vanilla, all three methods: 1) increase coding capacity (i.e., per input, each unit has a probability closer to 50% of being activated); 2) increase discrimination (classification accuracy); and 3) act as a regularizer (narrowing the range of activation levels for each unit). Interestingly, WeightNorm achieves lower accuracy and generates representations that are less homeostatic, compared to both BatchNorm and Synaptic Scaling (Figures 2 and 3). This suggests that learning algorithms are more efficient when coupled with homeostatic load balancing, and either without the other degrades performance. This perspective contributes to the growing list of explanations for why normalization is so useful in deep networks [3], [4], [5], [6], [8], and a natural next step is to develop a theoretical understanding for why stability (i.e., creating homeostatic representations) may actually promote plasticity (i.e., improving classification accuracy and learning efficiency), as opposed to being in conflict.

Moving forward, there are several challenges that remain in bridging the gap between normalization in the artificial and biological neural networks. First, the implementation details of both types of networks are well-acknowledged to be different [61]. For example, unlike most artificial networks, the brain has a strict division of excitatory and inhibitory neurons, which means different homeostasis rules can be applied to excitatory and inhibitory synapses [62]. Second, our model of synaptic scaling assumed that each hidden unit had the same target “fixed point”, whereas in reality, adjustable fixed points might further improve performance. Indeed, batch normalization allows the fixed points to be learned through the affine parameter, *β*. In artificial networks, fixed points could vary based on the dataset, network architecture, or other hyper-parameters. In the brain, different cell types may use different fixed points, or fixed points of a single cell may change during different phases of training. Third, it is unclear how the time-scales of homeostasis in the brain map to time-scales of learning in artificial networks. Normalization is typically applied per input or per batch in deep learning, but other time scales remain unexplored [35], [36], [63]. Similarly, normalization that operates simultaneously across different spatial scales (e.g., combining batch normalization and layer normalization) have also not been analyzed. Fourth, there are different constraints between what a hidden unit can store and compute and what a neuron can (likely) store and compute. For example, it seems plausible for a neuron to track its own mean firing rate over a given time window, but tracking its own variance seems trickier.

There are also several challenges in understanding the neuroscience of homeostasis that remain outstanding. For example, network-wide homeostasis, which goes beyond fixed points for individual neurons, has been observed in the brain, but the circuit mechanisms that give rise to these effects remain elusive. Further, it remains unclear what the advantages and disadvantages of different homeostatic mechanisms are, and when to use which. For example, many homeostatic plasticity mechanisms reset a neuron’s firing rate to a target firing rate on average; but when would it be appropriate to achieve this goal by modifying intrinsic excitability versus modifying pre- or post-synaptic weights? Indeed, there may be multiple means towards the same end, and it remains unclear what the trade-offs are among these different paths.

We hope these insights provide an avenue for building future collaborations, where computer scientists can use quantitative frameworks to evaluate how different plasticity mechanisms affect neural function. In return, neuroscientists can provide new perspectives on the benefits of normalization in neural networks and inspiration for designing new normalization algorithms based on neurobiological principles.

## Acknowledgments

The authors thank Sanjoy Dasgupta, Alexei Koulakov, Vishaal Krishnan, Ankit Patel, and Shyam Srinivasan for helpful comments on the manuscript. S.N. was supported by the Pew Charitable Trusts, the National Institutes of Health under Awards 1R01DC017695 and 1UF1NS111692, and funding from the Simons Center for Quantitative Biology at Cold Spring Harbor Laboratory. Y.S. was supported by a Swartz Foundation Fellowship.

## Notes

### Competing Interest Statement

The authors have declared no competing interest.

### Summary of Updates

updated acknowledgments.

